# Microbiome dysbiosis regulates the level of energy production under anaerobic condition

**DOI:** 10.1101/2021.05.11.443548

**Authors:** M. Shaminur Rahman, M. Nazmul Hoque, Joynob Akter Puspo, M. Rafiul Islam, Niloy Das, M. Anwar Siddique, M. Anwar Hossain, Munawar Sultana

## Abstract

The microbiome of the anaerobic digester (AD) regulates the level of energy production. To assess the microbiome dysbiosis in different stages of anaerobic digestion, we analyzed 16 samples dividing into four groups (Group-I = 2; Group-II = 5; Group-III = 5 and Group-IV = 4) through whole metagenome sequencing (WMS). The physicochemical analysis revealed that highest CH_4_ production (74.1%, on Day 35 of digestion) was associated with decreased amount of non-metal (phosphorus and sulfur) and heavy metals (chromium, lead and nickel). The WMS generated 380.04 million reads mapped to ~ 2800 distinct bacterial, archaeal and viral genomes through PathoScope (PS) and MG-RAST (MR) analyses. The PS analysis detected 768, 1421, 1819 and 1774 bacterial strains in Group-I, Group-II, Group-III and Group-IV, respectively which were represented by *Firmicutes*, *Bacteroidetes*, *Proteobacteria*, *Actinobacteria*, *Spirochaetes* and *Fibrobacteres* (> 93.0% of the total abundances). The archaeal fraction of the AD microbiomes was represented by 343 strains, of which 95.90% strains shared across these metagenomes. The indicator species analysis showed that *Methanosarcina vacuolate*, *Dehalococcoides mccartyi*, *Methanosarcina* sp. Kolksee and *Methanosarcina barkeri* were the highly specific for energy production in Group-III and Group-IV. However, most of the indicator phylotypes displayed reduced abundance in the initial stage of biogas production (Group-I and Group-II) compared to their increased relative abundances in Group-IV (Day 35). The correlation network analysis showed that different strains of *Euryarcheota* and *Firmicutes* phyla were associated with highest level (74.1%) of energy production (Group-IV). In addition to taxonomic dysbiosis, top CH_4_ producing microbiomes showed increased genomic functional activities related to one carbon and biotin metabolism, oxidative stress, proteolytic pathways, MT1-MMP pericellular network, acetyl-CoA production, motility and chemotaxis. This study reveals distinct changes in composition and diversity of the AD microbiomes including different indicator species, and their genomic features that are highly specific for energy production.

## Introduction

Bangladesh is experiencing rapidly increased energy consumption over the past two decades. Being one of the world’s most densely populated and least urbanized countries, around 72% of population of this country live in rural areas where there is no supply of natural gas, the main source of energy [1]. The access to clean and affordable energy is one of the prerequisites to achieve the sustainable development in rural areas. Upgrading existing biomass resources (i.e., animal manure, crop residues, kitchen and green wastes) to biogas shows significant promise in this respect [2]. The production of biogas through anaerobic digestion process cannot only provide fuel, but is also important for reduction of fertilizer nutrient utilization, rural forest conservation, protecting the environment, realizing agricultural recycling, as well as improving the sanitary conditions, in rural areas [3,4]. Biogas, produced from anaerobic digester (AD) is a is a relatively high-value fuel and continuous source of energy supply, and insurance of future energy in a sustainable manner [5]. Anaerobic environments play critical roles in the global carbon cycle through the digestion of organic agricultural waste, manure, municipal waste, digester materials, sewage, green waste or food. Biogas can significantly contribute to abate greenhouse gas emissions from livestock and agricultural farming at relatively lower mitigation costs. However, attention must be paid towards undesired emissions of methane (CH_4_) and nitrous oxide (N_2_O) from manure storage [6]. Moreover, anaerobic digestion of livestock manure improves organic fertilizer quality compared with undigested manure [7], and the load of the pathogenic microorganisms and related antimicrobial resistance is also decreased through the biological process of anaerobic degradation [8]. The rising energy prices and increasing concern of emission of greenhouse gases are the major concern for the people and agro-industries worldwide to consider the wider application of AD technology. This sustainable technology has been viewed as a way to address environmental concern through the generation of CH_4_ within engineered bioreactors, and thereby reducing the human dependence on fossil fuels [9,10]. The conversion of organic wastes, agricultural residues and renewable primary products into energy and other valuable products AD by the efficient process of the AD is also considered as a sustainable solution for resource recovery and renewable energy production underpinning the circular economy concept [9]. Furthermore, environment friendly renewable energy produced from locally available raw materials and recycled waste could thus contribute to climate change mitigation [5].

The renewable energy (CH_4_) produced from the anaerobic bioreactor (AD) which is independent of weather conditions could serve for the production of electricity, heat and fuels [11]. Anaerobic transformation of organic wastes in the AD is carried out by different bacterial and archaeal species, such as hydrolytic, acid forming, acetogenic, and methanogens which produce CO_2_ and CH_4_ as the main products of the digestion process [9]. Methane rich biogas (typically 50 – 70% methane, 30–50% CO_2_, with traces of H_2_S and other gases) is a clean, efficient, and renewable source of energy, which offers a multipurpose carrier of energy, and can be used as a substitute for other fuels [12]. Though biogas (CH_4_) is directly influenced by the composition of the AD microbiomes [9,13], the genomic potentials of the microbiomes favoring anaerobic metabolism to control the level of CH_4_ production is thermodynamically dependent on environmental parameters of the AD [14]. Diverse microbial communities are associated with biomass decomposition and CH_4_ production through the metabolic activities of substrate hydrolysis, acidogenesis, acetogenesis and methanogenesis [15]. The environmental and internal factors such as substrate ingredients, temperature, pH, level of CO_2_, O_2_ and H_2_S, and mixing or the geometry of the AD can be achieved through microbial selection or manipulation [9,13,16]. Therefore, a clear understanding of the structure, composition and diversity of the multifarious microbial community involved in biogas production is crucial for the optimization of their performance and stable operational process of the AD. Moreover, a detailed insight into relevant microbial metabolic pathways involved in CH_4_ synthesis and syntropy is essential to upsurge the yield of biogas. To reveal the dynamic changes of the exceedingly diverse and unified networks of AD microbiomes, few studies focused on the taxonomic and functional characterization of microbiomes originating from both laboratory-scale [9] and full-scale [9,17] biogas reactors under different prevailing ecological conditions [13,18]. Initially, AD microbiomes characterization mainly relied on conventional microbiology approaches. However, at the very beginning of the twentieth century more than 150 species of microorganisms have been identified from the anaerobic bioreactors through the application modern genomic approaches [19]. The current accelerated pace of genomic technology and the rapid incorporation of biotechnological techniques allowed us the rapid identification and characterization microorganisms such as *Clostridium bornimense*, *Herbinix hemicellulosilytica*, *Herbinix luporum*, *Herbivorax saccincola*, *Proteiniphilum saccharofermentans*, *Petrimonas mucosa*, *Fermentimonas caenicola*, and *Proteiniborus indolifex* or even their genomic features for the increased production of biogas [9,13,17,19–21]. The conventional culture-based techniques [22,23] for characterization of the microbiotas in different niches including controlled anaerobic chambers [11,24] has been replaced during the last decade by the rapid advances in high-throughput NGS technology and bioinformatics tools [25,26]. Despite, the 16S rRNA partial gene sequencing approach remained the most widely used genomic approach to study the microbiomes of the AD [12,27], several inherent limitations including the polymerase chain reaction (PCR) bias, inability to detect viruses, lower taxonomic resolution (up to genus level only), and limiting information on gene abundance and functional profiling have made this technique questionable [25,26]. Conversely, the whole metagenome sequencing (WMS) or shotgun approach which can identify the total microbial components of a sample (including viruses, bacteria, archaea, fungi, and protists) is being used prudently to decipher the phylogenetic composition, microbiome structure and diversity including profiling of their functional characteristics and interconnections [12,28].

To address the dynamic shifts in microbiome diversity and composition to be associated with different level of renewable energy production in the controlled AD, we present a comprehensive deep metagenomic (WMS) analysis of sixteen (n=16) samples collected from the same AD under different pH, CO_2_, O_2_ and H_2_S levels and temperature. Using a homogeneous mapping and annotation workflow associated with a de-replication strategy, our analyses identified ~ 2800 distinct bacterial and archaeal species along with their co-presence networking, antimicrobial resistance and metabolic functional profiling. This study therefore provides an opportunity to in-depth study the genetic potential and performance of microbial taxa represented by WMS, and to relate their activities to generate renewable energy under changing environmental conditions and process parameters.

## Materials and methods

### Digester setup and experiment design

The experiment was conducted using an anaerobic digester (AD) plant prepared with provisions to measure temperature, slurry and gas sample collection and substrate charging. The biogas plant consisted of a digester, inlet-chamber, three slurry outlet pipes, gas outlet pipe and thermometer (Fig S1). The AD that contains the substrate (organic wastes) and converts it into biogas and slurry was of 3000 L capacity, and made of flexible polyvinyl chloride (PVC) fabric with thickness: 1.2mm. There was a specialized ball valve which ensures the anaerobic condition within the digester and control the flow of substrates. The temperature of the AD was monitored using a probe. The experiment was conducted for 45 days (Day 0 to Day 44, July 15 to August 27, 2019). The AD was charged 14 times during first 36 days (Day 0-35) with 1,192 kg raw cow dung (highest input volume = 375 kg and lowest input volume = 35 kg) (Table S1). Initially, the digester was started with charge of 375 kg feedstock where the ratio of raw cow dung and active sludge was 1:1. The raw semi-solid cow dung (CD) was mixed with seed sludge from previous biogas plant (slurry) before charged into the AD. The AD was portable, light in weight, low cost and retains more heat inside.

### Sample collection and physicochemical parameters analysis

The representative samples (n=16) including CD, slurry and active sludge (AS) were collected and stored for subsequent analysis. The samples were categorized into four groups (Group-I, Group-II, Group-III and Group-IV) based on collection time (Day 0 to Day 35) and CH_4_ concentration. The samples of Group-I (n = 2) were collected at Day 0 (day of first input) when the CH_4_ concentration of the digesta was 0.0% with an average pH of 5.44. Likewise, Group-II samples (n = 5) were collected at Day 2 and Day 7 of the digestion process when the CH_4_ concentration and pH of the digesta were 21% and 5.44, and 34% and 5.56, respectively. The sampling of the Group-III (n = 5) was done at Day 10 and Day 27 of the digestion process having the CH_4_ concentration and pH of the digesta 47.4% and 5.97, and 58.2% and 6.87, respectively. The Group-IV included samples (n = 4) collected at Day 34 and Day 35 of digestion when the CH_4_ concentration and pH of the digesta were 71.4% and 6.99, and 74.1% and 7.01, respectively (Table 1). Therefore, highest CH_4_ production (74.1%) was recorded at Day 35 of the digestion process. In addition, data on physicochemical parameters were recorded up to Day 45 (Table 1). Total nitrogen (TN) content was measured by micro-Kjeldahl method [29] while phosphorus, potassium, heavy metals (Lead, Zinc, Nickel, Cadmium, Chromium), organic carbon, Sulphur and moisture content were determined by spectrophotometric molybdovanadate [30], flame photometric [31], atomic absorption spectrophotometric [30], wet oxidation, turbidimetric and gravimetric methods from the Department of Soil Science, University of Dhaka. The detection limit of metals was of the order of 0.1 μg L^−1^ [32].

**Table 1:** Metagenome samples and their groupings in the anaerobic digester according to experiment time and methane (CH4) concentration.

| Sample ID | Groups | Day | $\text{CH}_4$ (%) | pH | Purity<br>(280/260) | Concentration<br>(ng/$\mu\text{L}$) |
| --- | --- | --- | --- | --- | --- | --- |
| R1 | Group-I | 0 | 0 | 5.43 | 1.78 | 103.7 |
| M3 |  |  |  | 5.46 | 1.82 | 91.7 |
| O4 | Group-II | 2 | 21 | 5.44 | 1.88 | 61 |
| I5 |  |  |  |  | 1.88 | 77.7 |
| M6 |  |  |  |  | 1.87 | 73.4 |
| O7 |  | 7 | 34 | 5.56 | 1.87 | 46 |
| I9 |  |  |  |  | 1.88 | 87.2 |
| O10 | Group-III | 10 | 47.4 | 5.97 | 1.87 | 48.6 |
| M11 |  |  |  |  | 1.86 | 86.1 |
| I12 |  |  |  |  | 1.88 | 100.5 |
| O60 |  | 27 | 58.2 | 6.87 | 1.89 | 59.4 |
| O13 |  |  |  |  | 1.89 | 61.3 |
| O15 | Group-IV | 34 | 71.4 | 6.99 | 1.81 | 67.7 |
| I14 |  |  |  |  | 1.82 | 65.4 |
| O16 |  | 35 | 74.1 | 7.01 | 1.81 | 112.3 |
| I17 |  |  |  |  | 1.81 | 53.6 |

\*R1 = Raw cow dung, M3 = mixtures with raw cow dung and slurry from previous biogas plant as seed, O = Outlet samples, I = Input position samples, M = Middle position samples.

### Metagenomic DNA extraction, sequencing and bioinformatics analysis

We extracted the total genomic DNA (DNA) from 16 samples (Data S1) using an automated DNA extraction platform with DNeasy PowerSoil Kit (QIAGEN, Germany) according to manufacturer’s instructions. Extracted gDNA was quantified and purity checked through NanoDrop (ThermoFisher, USA) with an absorbance ratio of 260/280. Shotgun metagenomic (WMS) libraries were prepared with Nextera XT DNA Library Preparation Kit (Hoque et al., 2019), and paired-end (2×150 bp) sequencing was performed on a NovaSeq 6000 sequencer (Illumina Inc., USA) from the Macrogen Inc. (www.macrogen.com) Seoul, Republic of South Korea. The gDNA from sixteen samples generated 380.04 million reads, and the average reads per sample was 23.75 million (maximum = 24.79 million, minimum = 20.75 million) (Data S1). The low-quality reads from the generated FASTQ files were filtered and removed through BBDuk (with options k = 21, mink = 6, ktrim = r, ftm = 5, qtrim = rl, trimq = 20, minlen = 30, overwrite = true) (Stewart et al. 2018). In this study, on an average 21.45 million reads per sample (maximum=23.75 million, minimum=18.85 million) passed the quality control step (Data S1). Less than 100 hits were filtered for the downstream analysis. Read normalization in each sample was performed using median sequencing depth through Phyloseq (version 4.0) package in R.

### Microbiome analysis and AMR profiling

The taxonomic assignment of the generated WMS data was performed using both mapping based and open-source assembly-based hybrid methods of PahtoScope 2,0 (PS) [33] and MG-RAST 4.0 (MR) [34]. In PS analysis, the NCBI Reference Sequence Database (NCBI RefSeq Release 201) for bacteria and archaea was prepared by Kraken2 [35]. The reads were then aligned against the target (RefSeq) libraries using Minimap2 [36], and filtered to remove the reads aligned with the cattle genome (bosTau8) and human genome (hg38) using BWA [37] and samtools [38]. In PS analysis, we employed the PathoID module to get exact read counts for downstream analysis [33]. We simultaneously uploaded the raw sequences to the MR server with proper metadata. In MR analysis, the uploaded raw reads were subjected to optional quality filtering with dereplication and host DNA removal, and finally annotated for taxonomic assignment.

The within sample (alpha) diversity of microbial communities was calculated using the Shannon and Simpson diversity indices [39] through the “Vegan” package in R. To evaluate alpha diversity in different groups, we performed the non-parametric test Kruskal-Wallis rank-sum test. The diversity across the sample groups (Beta-diversity) was measured with the principal coordinate analysis (PCoA) using Bray–Curtis dissimilarity matrices, and permutational multivariate analysis of variance (PERMANOVA) with 999 permutations to estimate a p-value for differences among groups [40]. Phyloseq and Vegan packages were employed for these statistical analyses [41]. Indicator species specific to a given sample group (having ≥ 1000 reads assigned to a taxon) were identified based on the normalized abundances of species using the R package indicspecies [42], and the significant indicator value (IV) index was calculated by the 999-permutation test. Larger IV indicates greater specificity of taxa and p < 0.05 was considered statistically significant [42]. Network analysis was used to explore the co-occurrence patterns of the energy producing bacterial and archaeal taxa across the metagenome groups. In addition, the Spearman’s correlation coefficient and significance tests were performed using the R package Hmisc. A correlation network was constructed and visualized with Gephi (ver. 0.9.2).

We detected the antibiotics resistance genes (ARGs) among the microbiomes of four metagenomes through the ResFinder 4.0 database (https://doi.org/10.1093/jac/dkaa345). The ResFinder database was integrated within AMR++ pipeline [43] to identify the respective genes and/or protein families [28]. In addition, the OmicCircos (version 3.9) [28], an R package was used for circular visualization of both diversity and composition of ARGs across the metagenomes under study.

### Functional profiling of the microbiomes

In addition to taxonomic annotations, the WMS reads were also mapped onto the Kyoto Encyclopedia of Genes and Genomes (KEGG) database [44], and SEED subsystem identifiers, respectively, on the MR server [34] for metabolic functional profiling. The functional mapping was performed with the partially modified set parameters (*e-*value cutoff: 1×10^−30^, min. % identity cutoff: 60%, and min. alignment length cutoff: 20) of the MR server [28].

### Statistical analysis

To evaluate differences in the relative percent abundance of taxa in AD (Group-I, Group-II, Group-III and Group-IV) for PS data, we used the non-parametric test Kruskal-Wallis rank sum test. We normalized the gene counts by dividing the number of gene hits to individual taxa/function by total number of gene hits in each metagenome dataset to remove bias due to differences in sequencing efforts. The non-parametric test Kruskal-Wallis rank sum tests were also performed to identify the differentially abundant SEED or KEGG functions (at different levels), and antimicrobial resistance (ARGs) in four metagenomes. All the statistical tests were carried out using IBM SPSS (SPSS, Version 23.0, IBM Corp., NY USA). To calculate the significance of variability patterns of the microbiomes (generated between sample categories), we performed PERMANOVA (Wilcoxon rank sum test using vegan 2.5.1 package of R 3.4.2) on all four sample types at the same time and compared them pairwise. A significance level of alpha = 0.05 was used for all tests.

## Results

### Physicochemical properties of substrate and digesta

The physicochemical properties of the digester feedstock before and after the anaerobic digestion of cow dung are shown in Figure 1 and Table 2. The fermentation was run for 44 days. Periodic increments of the organic loading rate (OLR) resulted in increased biogas production (Fig 1A, Table S1). This trend was observed until Day 35, after which biogas production began to decline gradually, although the OLR was kept constant. The log phase of methane (CH_4_) production started around the second day of the experiment, and reached its maximum percentage (74.1%) at Day 35 of the digestion process. It should be pointed out that after Day 35, the CH_4_ percentage started to decrease, reaching 59.2% on Day 44 (Table 1). Average concentration (%) CO_2_ was observed 39.52 (minimum = 27.7, maximum = 56) throughout 44 days of the digestion process. Concentration of H_2_S was maximum (938 ppm) at Day 3, and later on the concentration fluctuated based on feeding (Fig 1A, Table S1). The overall environmental temperature, AD temperature, AD pressure and humidity were 34.75 °C (maximum = 38.8 °C, minimum = 32.0 °C), 34.46 °C (maximum = 51.0 °C, minimum = 0.0 °C), 22.52 mb (maximum = 56.41 mb, minimum = 0.0 mb), and 55.5 % (maximum = 94.0 %, minimum = 42.0 %) (Fig 1B, Table S2). On Day 35 of the digestion, when maximum methanogenesis was observed, the concentration of organic carbon (OC) and total nitrogen (TN) in the fermentation pulp were 15.48% and 1.22%, respectively, whereas the concentration of OC and TN in the slurry (CD + seed sludge; Day-0) were 34.39% and 1.96%, respectively (Table 2). The overall C/N ratio of the feedstock also gradually decreased with the advent of anaerobic digestion process, and found lowest (12.7:1) at Day 35. Similarly, the amount of non-metallic element (phosphorus and sulfur) and heavy metals (chromium, lead and nickel) content significantly decreased at the Day 35 of the digestion process (Table 2). However, the amount of zinc and copper did not vary significantly throughout the digestion period (Table 2).

**Table 2:** Physicochemical properties of raw cow dung, slurry and active sludges.

| Parameters | Cow dung (CD; Day-0) | Slurry (CD + seed sludge; Day-0) | Active sludge (AS; Day-35, highest CH<sub>4</sub> concentration) |
| --- | --- | --- | --- |
| Moisture (%) | 23.32 | 86.16 | 90.03 |
| Organic carbon (%) | 34.39 | 36.83 | 15.48 |
| Total nitrogen (%) | 1.82 | 1.96 | 1.22 |
| C: N | 18.9:1 | 18.8:1 | 12.7:1 |
| Phosphorus (%) | 0.249 | 0.461 | 0.070 |
| Sulphur (%) | 0.522 | 0.579 | 0.038 |
| Zinc (mg/kg) | 16.24 | 17.56 | 15.58 |
| Copper (mg/kg) | 2.88 | 3.59 | 2.13 |
| Chromium (mg/kg) | 11.97 | 11.33 | 0.23 |
| Cadmium (mg/kg) | BDL | BDL | 0.04 |
| Lead (mg/kg) | 3.44 | 3.26 | 1.06 |
| Nickel (mg/kg) | 3.28 | 6.70 | 0.34 |

CD: raw semi-solid cow dung which was mixed with water; Slurry: mixture of CD and seed sludge from previous biogas plant; AS: slurry from the AD when the gas production rate was the highest (i.e. 74.1% CH<sub>4</sub> concentration); and BDL: below detection limit.

**Fig 1:**
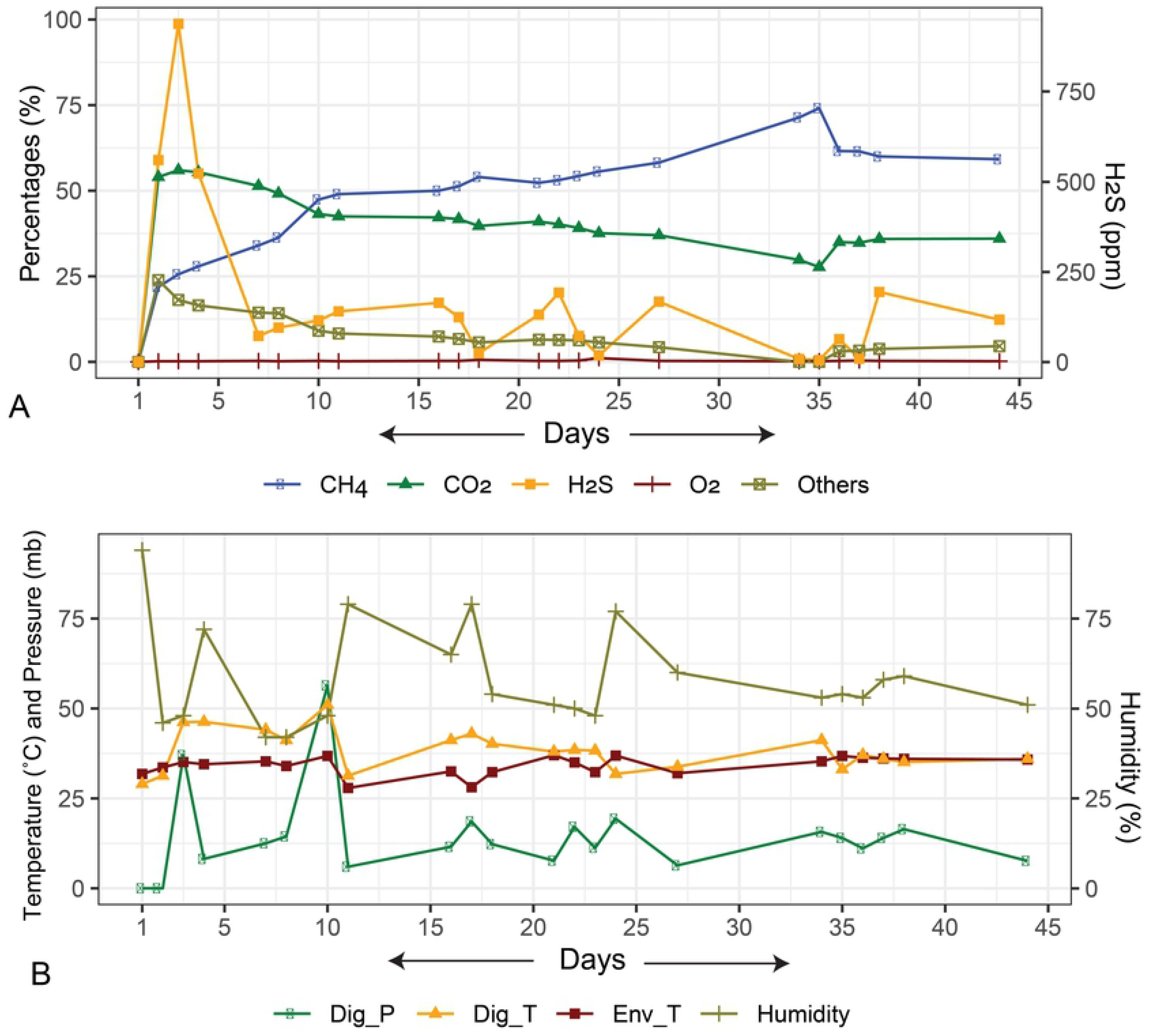
Dynamic changes in the physicochemical parameters of the anaerobic digester (AD) over the study period.

### Microbiome composition and diversity in anaerobic digester

The whole metagenome sequencing (WMS) of 16 sample libraries resulted in 380.04 million reads passing 343.26 million reads quality filters, which corresponded to 90.32% total reads (individual reads per sample are shown in Data S1). The major microbial domain in all samples was Bacteria with an abundance of 81.80%, followed by archaea (15.43%), and viruses (2.77%) (Data S1).

The alpha diversity (i.e., within-sample diversity) of the AD microbiomes was computed using the Shannon and Simpson estimated indices (i.e., a diversity index accounting both evenness and richness) at the strain level. In this study, both Shannon and Simpson indices estimated diversity significantly varied across the four sample groups (p = 0.03541, Kruskal-Wallis test). The pair-wise comparison of the within sample diversity revealed that the microbiomes of the Group-II significantly differed with those of Group-III and Group-IV (p= 0.048, Wilcoxon rank sum test for each) compared to Group-I (p= 0.91, Wilcoxon rank sum test) (Fig 2A, B). The rarefaction analysis of the observed species showed a plateau after, on average, 21.45 million reads (Fig S2, Data S1)-indicating that the coverage depth for most samples was sufficient to capture the entire microbial diversity. We also observed significant differences in the microbial community structure among the four metagenome groups (i.e., beta diversity analysis). Principal coordinate analysis (PCoA) at the strain level (Fig 2C), showed a distinct separation of samples by the experimental groups. Besides, we found significant (p = 0.032, Kruskal Wallis test) differences in the abundance of ARGs and metabolic functional genes/pathways (Data S2) which could strongly modulate the level of energy production through microbiome dysbiosis in the AD.

**Fig 2:**
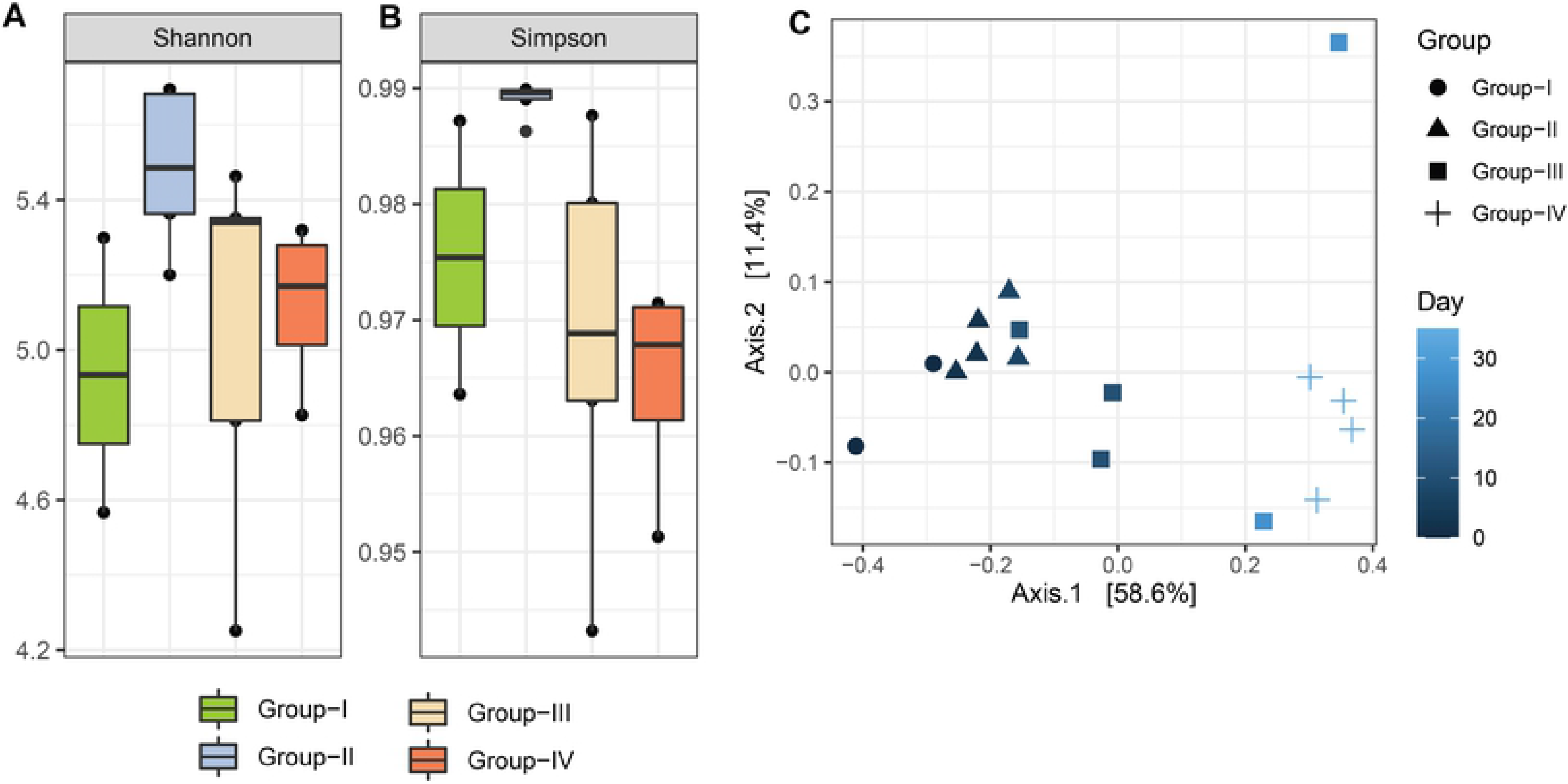
Microbiome diversity in four metagenomic groups of the anaerobic digestate samples. Box plots showing significant differences in observed species richness in AD associated microbiome. (A-B) Alpha diversity, as measured by PathoScope (PS) analysis using the Shannon and Simpson diversity indices, revealed distinct microbiome diversity across four metagenome samples (p = 0.03541, Kruskal-Wallis test). (C) The experimental groups were clearly separated by principal coordinate analysis (PCoA), which was measured using non-metric multidimensional scaling (NMDS) ordination plots. The different shapes represent the assigned populations in four metagenomes. As the day progresses, the group color becomes lighter. Values in parentheses represent the fraction of the total variance explained by each PCoA axis.

In this study, on an average 0.43% WMS reads (assigned for r RNA genes) mapped to 28, 110 and 552 bacterial phyla, orders and genera respectively, and relative abundance of the microbiome differed significantly (p = 0.034, Kruskal-Wallis test) across the metagenome groups (Data S1). We observed significant shifts/dysbiosis in the microbiome composition at strain level. The PS analysis detected 2,513 bacterial strains across the four metagenomes, of which 768, 1421, 1819 and 1774 strains were found in Group-I, Group-II, Group-III and Group-IV metagenomes, respectively. Only, 18.34% detected strains were found to be shared across the four energy producing metagenomes (Fig 3, Data S1). The archaeal fraction of the AD microbiomes was represented by 5, 17, 61 and 343 archaeal phyla, orders, genera and strains, respectively, and the relative abundance of these microbial taxa also varied significantly among the four metagenome groups. Remarkably, 95.90% (329/343) of the detected archaeal strains shared across these metagenomes (Fig 3, Data S1). In addition, 472, 536, 535 and 536 strains of bacterial viruses (bacteriophages) were identified in Group-I, Group-II, Group-III and Group=IV metagenomes, respectively.

**Fig 3:**
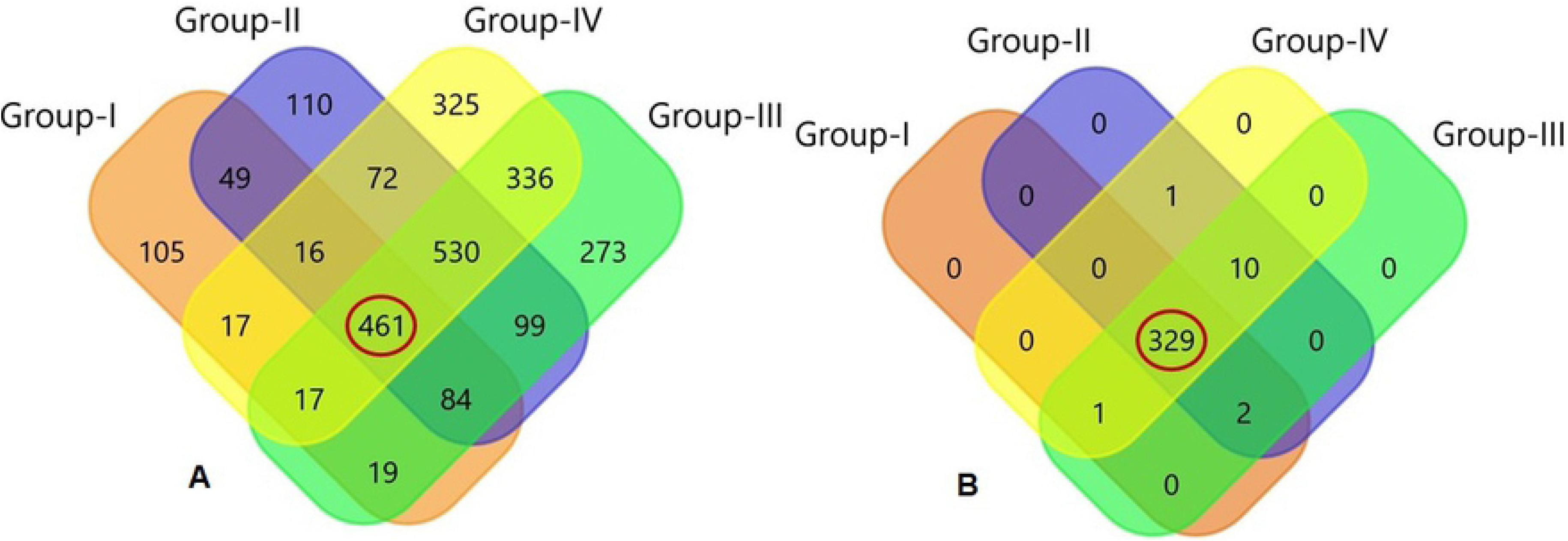
Unique and shared taxonomic composition of AD associated microbiome. Four metagenomics samples were represented by Venn diagrams depicting the core unique and shared microbiomes. (A) Venn diagram showing unique and shared bacterial strain by PS analysis, (B) Venn diagram showing unique and shared archaeal strain by PathoScope analysis. Microbiome sharing between the conditions are indicated by red circles. More information on the taxonomic result is also available in Data S1.

### Microbial community dynamically changed over time in the anaerobic digester

Significant changes in the abundances of core microbial groups were observed under anaerobic condition of the AD. At phylum level, the AD metagenome was dominated by *Firmicutes*, *Bacteroidetes*, *Proteobacteria*, *Actinobacteria*, *Spirochaetes* and *Fibrobacteres* comprising > 93.0% of the total bacterial abundances. Among these phyla, *Firmicutes* was the most abundant phylum with a relative abundance of 41.94%, 37.99%, 40.40% and 38.96% in Group-1, Group-II, Group-III and Group-IV, respectively. The relative abundance of *Bacteroidetes* (from 37.87% in Group-I to 22.40% in Group-IV) and *Actinobacteria* (from 3.94% in Group-I to 3.30% in Group-IV) gradually decreased with the advance of AD digestion time. Conversely, relative abundance of *Proteobacteria* (from 8.08% in Group-I to 18.92% in Group-IV) and *Spirochaetes* (from 1.28% in Group-I to 3.70% in Group-IV) gradually increased with the increase of anaerobic digestion time in AD. The rest of phyla also differed significantly across these four groups keeping comparatively higher relative abundances during highest CH_4_ producing stage (Group-IV) of the AD. Similarly, *Clostridiales* and *Bacteroidales* were identified as the top abundant order in Group-1, Group-II, Group-III and Group-IV with a relative abundance of 32.37%, 27.81%, 29.22% and 27.87%, and 32.49%, 27.42%, 27.53% and 14.94%, respectively (Data S1).

The structure and relative abundances of the bacteria at the genus level also showed significant differences (p = 0.031, Kruskal-Wallis test) across the study groups. In Group-I, Group-II and Group-III metagenomes, *Bacteroides* was the most abundant bacteria with a relative abundance of 18.10%, 14.90% and 15.16%, respectively, but remained lower (8.31%%) in Group-IV samples. *Clostridium* was found as the second most predominant bacterial genus, and the relative abundance of this bacterium was 11.92%, 11.13%, 11.73% and 12.15% in Group-1, Group-II, Group-III and Group-IV, respectively. The relative abundance of *Ruminococcus*, *Eubacterium*, *Parabacteroides*, *Fibrobacter*, *Paludibacter*, *Porphyromonas* and *Bifidobacterium* gradually decreased with the increase of energy (CH_4_) production rate, and remained lowest in Group-IV. Conversely, *Candidatus*, *Bacillus*, *Treponema* and *Geobacter* showed an increasing trend in their relative abundances gradually with the advance of digestion time and remained lowest in relative abundances in Group-IV. The rest of the bacterial genera had lower relative abundances in four metagenomes of the AD (Fig S3, Data S1).

In this study, *Methanosarcina* was the most abundant archaeal genus, and the relative abundance of this genus remained two-fold higher in Group-III (35.84%) and Group-IV (36.53%) compared to Group-I (17.52%) and Group-II (18.32%). Notably, the relative abundance of *Methanoculleus* was found higher in Group-II (11.59%) and Group-IV (13.80%) and lowest in Group-I (3.46%). Likewise, *Methanobrevibacter* was predominantly abundant at the initial phage of digestion (highest in Group-I; 19.35%) and remained lowest in abundance in the top CH_4_ producing metagenome (Group-IV; 5.01%). Besides these genera, *Methanothermobacter* (5.30%), *Methanosaeta* (5.16%), *Methanococcus* (4.74%), *Thermococcus* (2.96%), *Methanocaldococcus* (2.53%), *Pyrococcus* (2.35%), *Methanosphaera* (2.32%) *Methanococcoides* (2.10%) and *Archaeoglobus* (2.01%) were the predominantly abundant archaeal genera in Group-I samples and their relative abundances gradually decreased with the increase of energy production (Fig S4, Data S1). On the other hand, *Methanoregula* (6.43%), *Methanosphaerula* (2.99%), *Methanoplanus* (2.37%) and *Methanohalophilus* (1.39%) were the most abundant archaeal genera in Group-IV metagenome. The rest of the genera remained much lower (< 1.0%) in relative abundances but varied significantly across the four metagenomes (Fig S4, Data S1).

The strain-level composition, diversity and relative abundances of the microbiomes across four metagenomes revealed significant variations (p = 0.011, Kruskal-Wallis test) (Fig 4, Data S1). In this study, 2,513 bacterial and 343 archaeal strains were detected, of which 18.35% (461/2513) bacterial and 95.92% archaeal strains shared across the study metagenomes (Fig 3, Data S1). Most of the bacterial strains detected were represented by the phylum *Firmicutes* followed by *Bacteroidetes*, *Gammaproteobacteria* and *Betaproteobacteria* (Fig S5, Data S1). Of the detected strains, methanogenic archaeal strains were more prevalent (higher relative abundances) compared to bacterial strains, and this stain-level microbiome profiling was more evident in highest energy producing metagenome group (Group-IV). The most prevalent energy producing archaeal strains in Group-IV were *Methanosarcina vacuolata* Z-761 (17.31%), *Methanosarcina* sp. Kolksee (16.63%), *Methanoculleus marisnigri* JR1 (5.0%), *Methanothrix soehngenii* GP6 (4.61%), *Methanobacterium formicicum* DSM 1535 (3.60%), *Methanoculleus* sp. MAB1 (2.07%) and *Methanoculleus bourgensis* DSM 3045 (2.07%), and rest of strains had lower (< 2.0%) relative abundances (Fig 4, Data S1). Moreover, the relative abundances of these strains gradually increased with the increase of energy production (lowest relative abundance in Group-I and highest relative abundance in Group-IV) (Data S1). Conversely, the relative abundances of most of the bacterial strains identified gradually decreased with the advance of digestion time (increase of energy production), and mostly remained higher in relative abundances in Group-I (Data S1). Of the top abundant bacterial strains, *Bifidobacterium pseudolongum* subsp. globosum DSM 20092 (12.0%), *Phocaeicola dorei* DSM 17855 (6.61%), *Fibrobacter succinogenes* subsp. succinogenes S85 (4.57%), *Faecalibacterium prausnitzii* M21/2 (2.89%), *Clostridiales bacterium* CCNA10 (2.78%), and *Flintibacter* sp. KGMB00164 (2.07%) were found in Group-I, and their abundances gradually decreased with the increase of level of CH_4_ production. In addition, *Dysosmobacter welbionis* J115 (5.48%) remained more prevalent in Group-II (Fig 4, Data S1). The rest of the bacterial strains were less abundant (<2.0%) across the four metagenomes (Fig 4, Data S1). The viral fraction of the microbiomes mostly dominated by different strains of bacteriophages such as *Gordonia* phage Secretariat (16.12%), *Streptomyces* phage Bing (5.33%) and *Arthrobacter* phage Gordon (5.05%) in Group-I, *Megavirus chiliensis* (1.81%), *Acanthamoeba polyphaga* moumouvirus (1.60%) and *Orpheovirus* IHUMI-LCC2 (1.50%) in Group-II, *Stenotrophomonas* phage Mendera (4.88%), *Choristoneura fumiferana* granulovirus (3.0%) and *Gordonia* phage Secretariat (2.55%) in Group-III and *Stenotrophomonas* phage Mendera (2.58%), *Choristoneura fumiferana* granulovirus (2.37%) and *Bacillus* phage Mater (1.47%) in Group-IV (Data S1).

**Fig 4:**
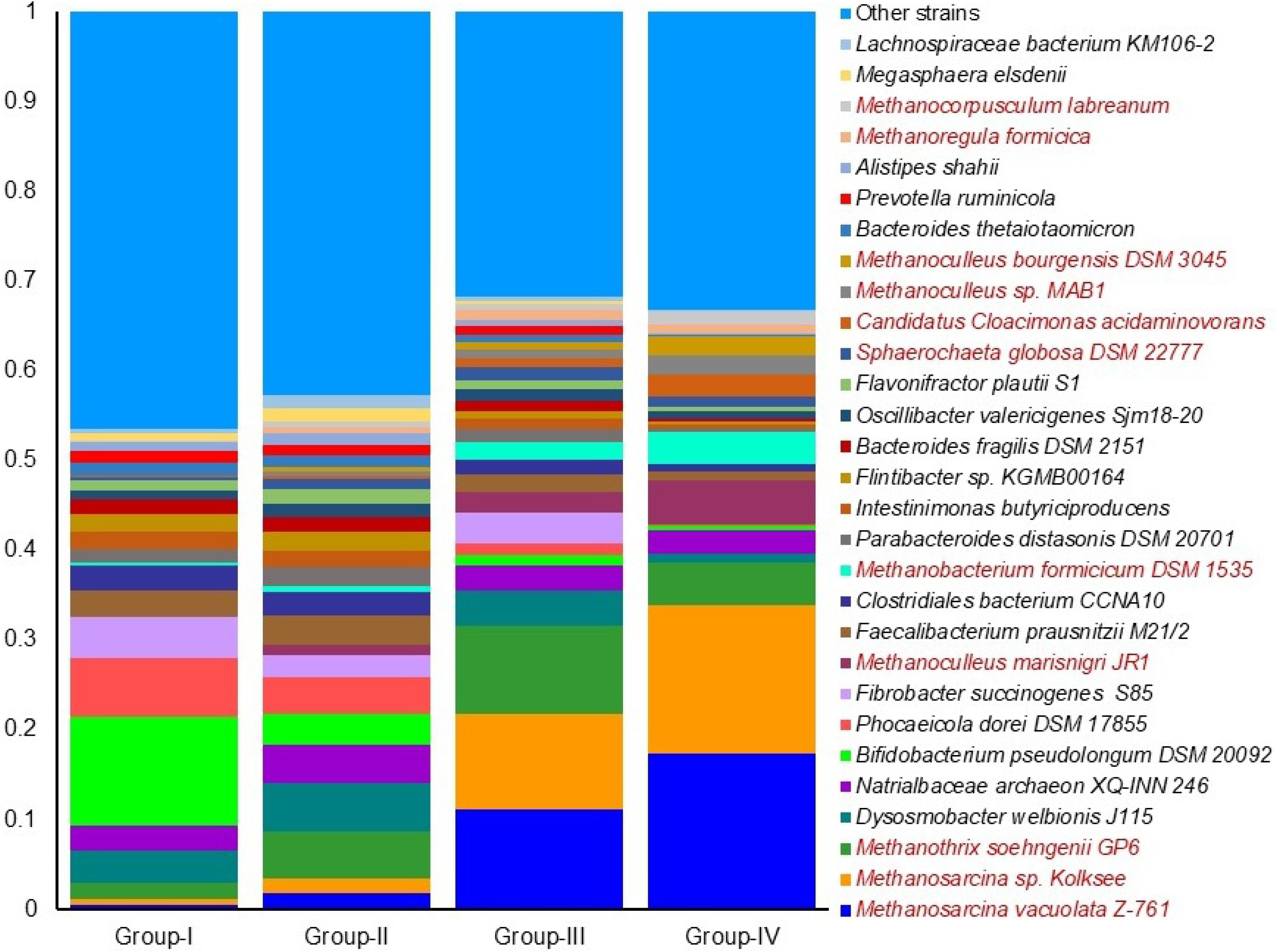
The strain level taxonomic abundance of anaerobic digestion driving microbiome. Stacked bar plots showing the relative abundance and distribution of the 30 most abundant strains, with ranks ordered from bottom to top by their increasing proportion among the four metagenomics groups. Only the 29 most abundant strains are shown in the legend, with the remaining strains grouped as ‘Other strains’. Each stacked bar plot represents the abundance of bacteria in each sample of the corresponding category. The relative abundances of archaeal strains (red colored) steadily improved as energy demand increased (lowest relative abundance in Group-I and highest relative abundance in Group-IV). In contrast, the relative abundances of most of the known bacterial strains gradually decreased with the passage of time (increased energy production), and mostly remained higher in Group-I and lower in Group-IV.

### Identification of potential indicator species and their co-occurrence

To identify microbial taxa (bacteria and archaea) that could discriminate across the four metagenome groups of the AD in terms of energy production (% CH_4_), the indicator species analysis (ISA) was performed both in individual group and combination basis, as shown in (Fig 5). Indicator species were those which were significantly more abundant and present in all samples belonging to one group, and also absent or low abundance in the other group (Fig 5, Data S1). The core taxa were selected based on their relative frequency (≥75% occurrence in each of the four groups) (Data S1). Although, 26, 3 and 19 indicator species were found in Group-I, Group-II and Group-IV, respectively, and no indicator species were identified in Group-III (Fig 5A, Data S1). Higher indicator values (IVs) suggested better performances in the microbial signature of the assigned taxa. *Desulfosporosinus youngiae*, *Treponema caldarium*, *Pseudoclostridium* thermosuccinogenes, Dehalobacterium formicoaceticum, Methanofollis liminatans, Methanoregula boonei, Syntrophomonas wolfei, Hungateiclostridium clariflavum, Candidatus Cloacimonas acidaminovorans and Methanocorpusculum labreanum were highly specific for energy production in Group-IV (highest CH_4_ production rate; 74.1%), with IVs of 0.983, 0.978, 0.949, 0.907, 0.887, 0.885, 0.882, 0.851, 0.795 and 0.786, respectively (Fig 5A; Data S1). Considering the combined group effects of the indicator species associated with energy production, our analysis revealed that *Methanosarcina vacuolate*, *Dehalococcoides mccartyi*, *Methanosarcina* sp. Kolksee and *Methanosarcina barkeri* in Group-III + Group-IV (top CH_4_ producing groups) having IVs of 0.88, 0.887, 0.879 and 0.879, respectively were highly specific for energy production (Fig 5B, Data S1). All of the indicator phylotypes displayed reduced abundance in the initial stage of biogas production (Group-I and Group-II, lower CH_4_ production rate) compared to their increased relative abundance up to Day 35 of the experiment (in Group-III and Group-IV) (Data S1).

**Fig 5:**
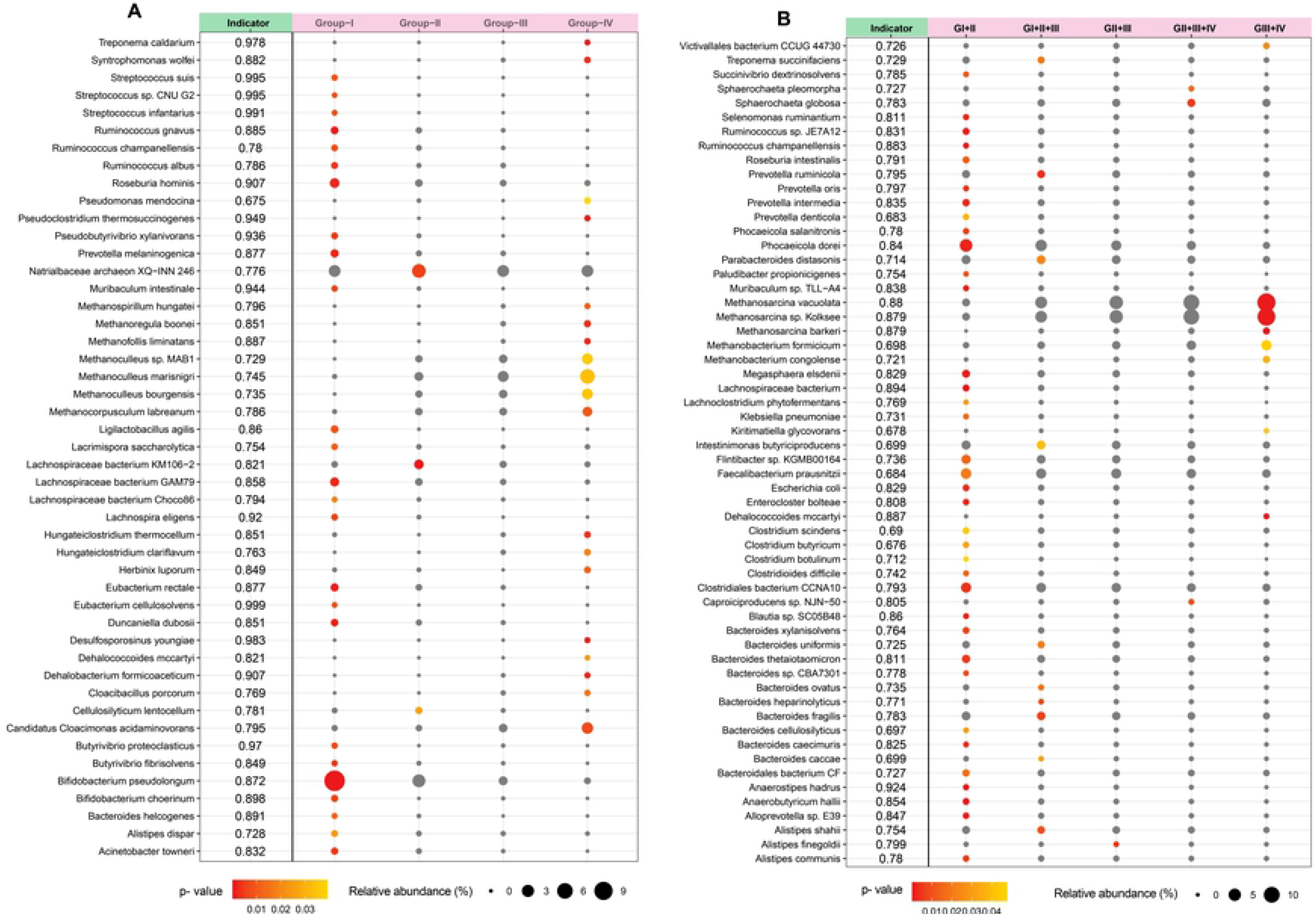
Indicator species analysis of AD microbiome within four metagenomics groups. (A) Individual group effects of the indicator species associated with energy production, (B) combined group effects of the indicator species associated with energy production. Indicator values (IndVal) are shown next to the taxonomic information for the indicator taxa as indicated by Indicator. Size of symbol is proportional to the mean relative abundance in that group of AD. Red symbols indicate for which group the taxon is an indicator. Gray symbols indicate group that contain a taxon, but for which that taxon is not an indicator taxa. Higher indicator values (IVs) suggested better performances in the microbial signature of the assigned taxa.

We then visualized networks within each metagenome group of the AD for both positive and negative co-occurrence relationships (Fig 6, Data S1). The correlation networks analysis was performed based on the significantly altered species (n=106) in different groups as revealed by indispecies analysis. This network analysis explored significant association (p = 0.021, Kruskal-Wallis test) in the co-occurrence patterns of the energy producing microbial taxa (species and/or strains) based on their relative abundances in four metagenome groups. In the correlation network of four metagenomes; Group-I to Group-IV), *Firmicutes* and *Bacteroidetes* exhibited strongest relation. The resultant network consists of 106 nodes (17 in Group-I, 58 in Group-II, 5 in Group-III and 26 in Group-IV) which were clearly separated into four modules/clusters (Fig 6). Taxa in the same group may co-occur under the same AD conditions (temperature, O_2_ and H_2_S percentage, pressure and humidity). Across different metagenome groups of AD, *Firmicutes*, *Bacteroidetes*, *Actinobacteria* and *Proteobacteria* were the top abundant phyla in Group-I and Group-II with a cutoff of 1.0 while *Bacteroidetes* and *Chlorhexi* in Group-III, and *Euryarcheota* and *Firmicutes* in Group-IV were designated as the top abundant phyla with a cutoff of 1.0 (Fig 6). However, when moving down to the species-level in microbiome co-occurrence in the AD, keystone taxa were much more consistent between networks with different correlation cutoffs. These results reveal that applying the same conditions in the AD for energy production, network elements must happen under careful consideration of the parameters used to delineate co-occurrence relationships. The positive correlations between Group-I and Group-II were observed among the microbiomes of the AD while Group-III and Group-IV showed negative correlations in terms of energy production with the microbial taxa of other two groups (Fig 6). These findings therefore suggest that different strains of *Euryarcheota* and *Firmicutes* phyla were negatively correlated but associated with highest level of energy production (highest % of CH_4_; Group-IV).

**Fig 6:**
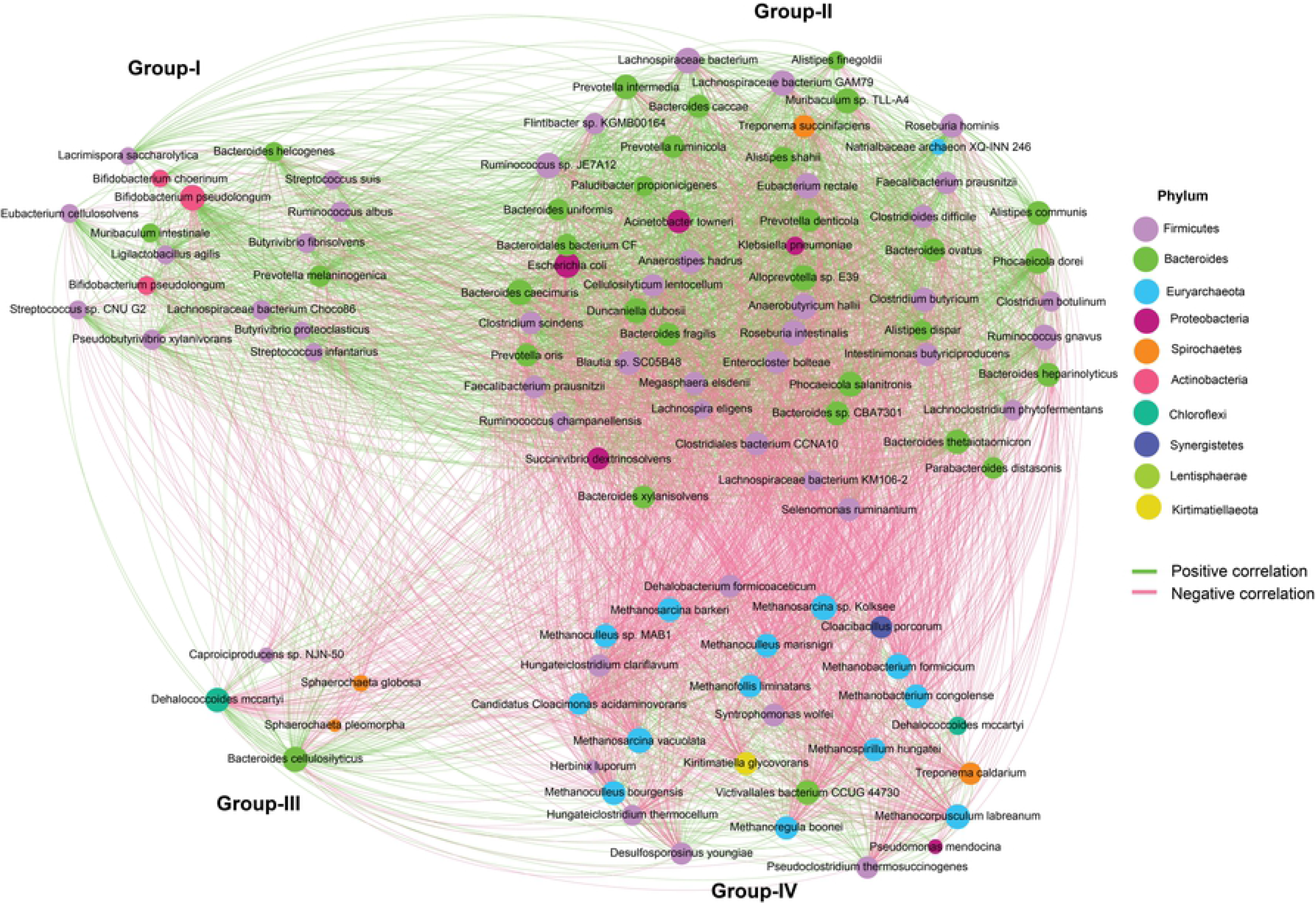
Microbiome co-occurrence in the AD within the four metagenomic groups. The microbiomes of the AD showed positive associations between Groups I and II, while Groups III and IV showed negative correlations in terms of energy production with the microbial taxa of the other two groups. Nodes are colored by taxonomy with labelled genera names. The positive correlation is represented by the green line, while the negative correlation is represented by the red line.

### Genomic functional potentials of the anaerobic microbiomes

In this study, there was a broad variation in the diversity and composition of the antimicrobial resistance genes (ARGs) (Fig 7, Data S2). The results of the present study revealed significant correlation (p = 0.0411, Kruskal-Wallis test) between the relative abundances of the detected ARGs and the relative abundance of the associated bacteria found in four metagenomes (Data S2). ResFinder identified 49 ARGs belonged to 19 antibiotic classes distributed in 2,513 bacterial strains (Data S2). The Group-III microbiomes harbored the highest number of ARGs (42), followed by Group-II (38), Group-IV (29) and Group-I (22) microbes (Fig 7, Data S2). The tetracyclines (doxycycline and tetracycline) resistant gene, *tet*Q had the highest relative abundance (23.81%) in Group-I associated bacteria followed by Group-II (22.85%), Group-III (16.49%) and Group-IV (6.73%)–microbes. Macrolides (erythromycin and streptogramin B) resistant genes such as *mef*A (16.80%), *mef*B (15.32%) and *msr*D (11.10%) had higher relative abundances in highest CH_4_ producing metagenome compared to other metagenome groups. The broad-spectrum beta-lactams resistant gene, *cfx*A2-6 was found as the common ARG among the microbiomes of four metagenomes, displaying the highest relative abundance (35.58%) in inoculum (Group-I) microbiota followed by Group-II (23.09%), Group-III (8.02%) and Group-IV (0.14%) microbiomes. The rest of the ARGs also varied in their expression levels across the four metagenomes, being more prevalent in the Group-III microbiomes (Fig 7). In addition to these ARGs, the highest CH_4_ producing microbiomes were enriched with the higher relative abundance of genes coding for cobalt-zinc-cadmium resistance (18.85%), resistance to chromium compounds (12.17%), arsenic (6.29%), zinc (4.96%) and cadmium (3.26%) resistance compared to the microbes of other three metagenomes (Fig S6, Data S2). By comparing the possible mechanisms of the detected ARGs, we found that antibiotic efflux pumps associated resistance had the highest level of expression in the anaerobic microbiomes of the AD followed by antibiotic inactivation, enzymatic inactivation and modification, antibiotic target protection/alteration, and folate pathway antagonist-attributed resistance mechanisms (Fig S6, Data S2).

**Fig 7:**
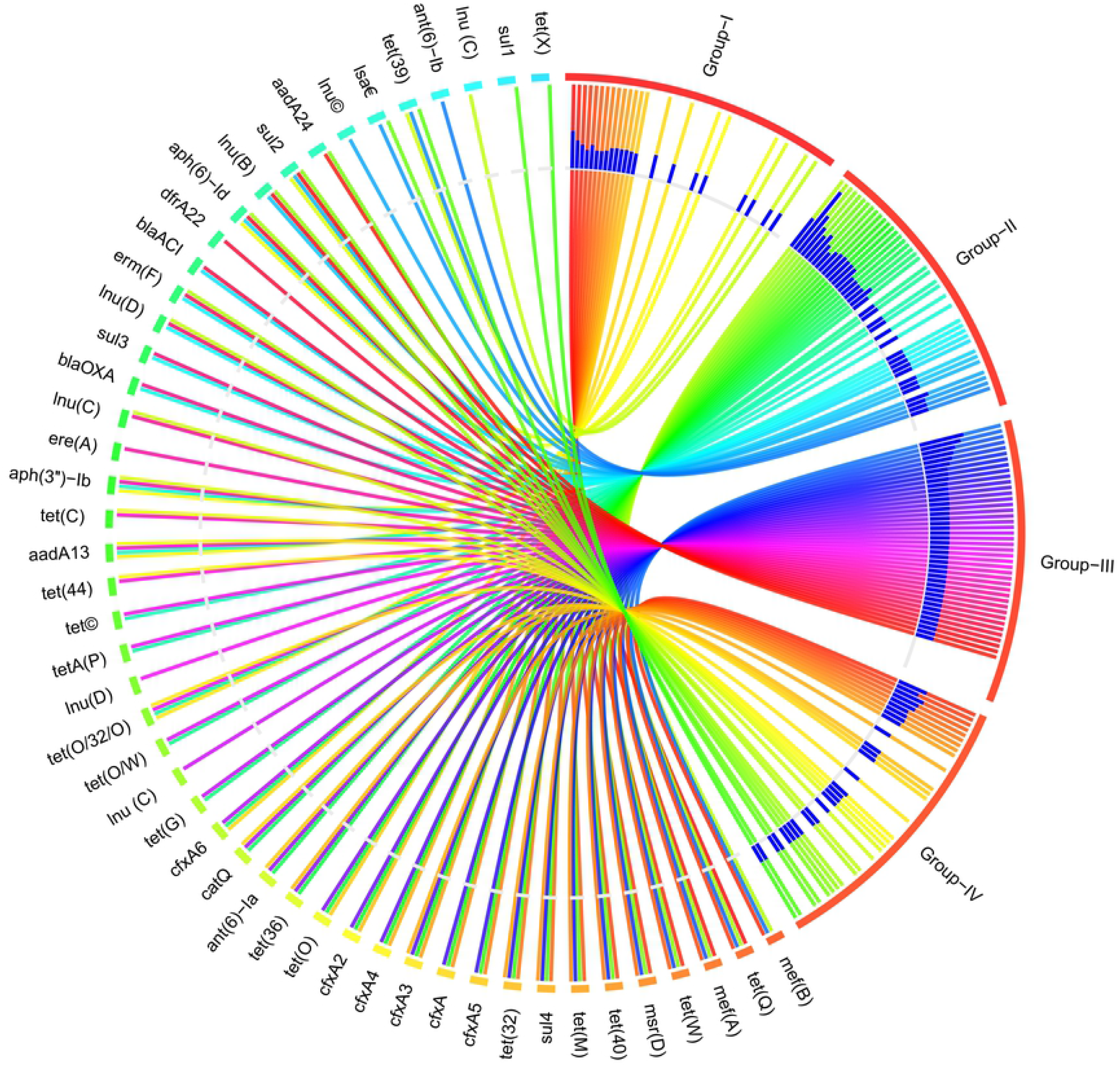
Antibiotics resistance genes (ARGs) detected in anaerobic digestion driving microbiome. The circular plot illustrates the distribution of 49 ARGs belonged to 19 antibiotic classes found across the four metagenomes. ARGs in the respective metagenome group are represented by different colored ribbons, and the inner blue bars indicate their respective relative abundances. Group-III associated microbiomes had the highest number of VFGs followed by Group-II, Group-IV and Group-I microbes.

Functional metabolic profiling of the gene families of the same KEGG pathway for AD microbiomes revealed significant differences (p = 0.012, Kruskal-Wallis test) in their relative abundances, and positive correlation with level of energy production. Of the detected KO modules, genes coding for CHO metabolism and genetic information and processing were top abundant, however did not vary significantly across the metagenome groups. Remarkably, the relative abundance of genes coding for energy metabolism, xenobiotics biodegradation and metabolism, butanoate metabolism, citrate synthase (*glt*A), succinyl-CoA synthetase subunits (*sucC*/D), pyruvate carboxylase subunits(*pycA*) and nitrogen metabolism gradually increased with the increasing rate of CH_4_ production, and had several-fold over expression among the microbiomes of Group-IV. Conversely, fumarate hydratase (*fumA*/B), malate dehydrogenase (*mdh*) and bacterial secretion system associated genes were predominantly overexpressed in Group-I related microbiomes which gradually decreased with advance of digestion process, and remained more than two-fold lower expressed in the peak level of CH_4_ production (lowest in Group-IV) (Fig 8A, Data S2).

**Fig 8:**
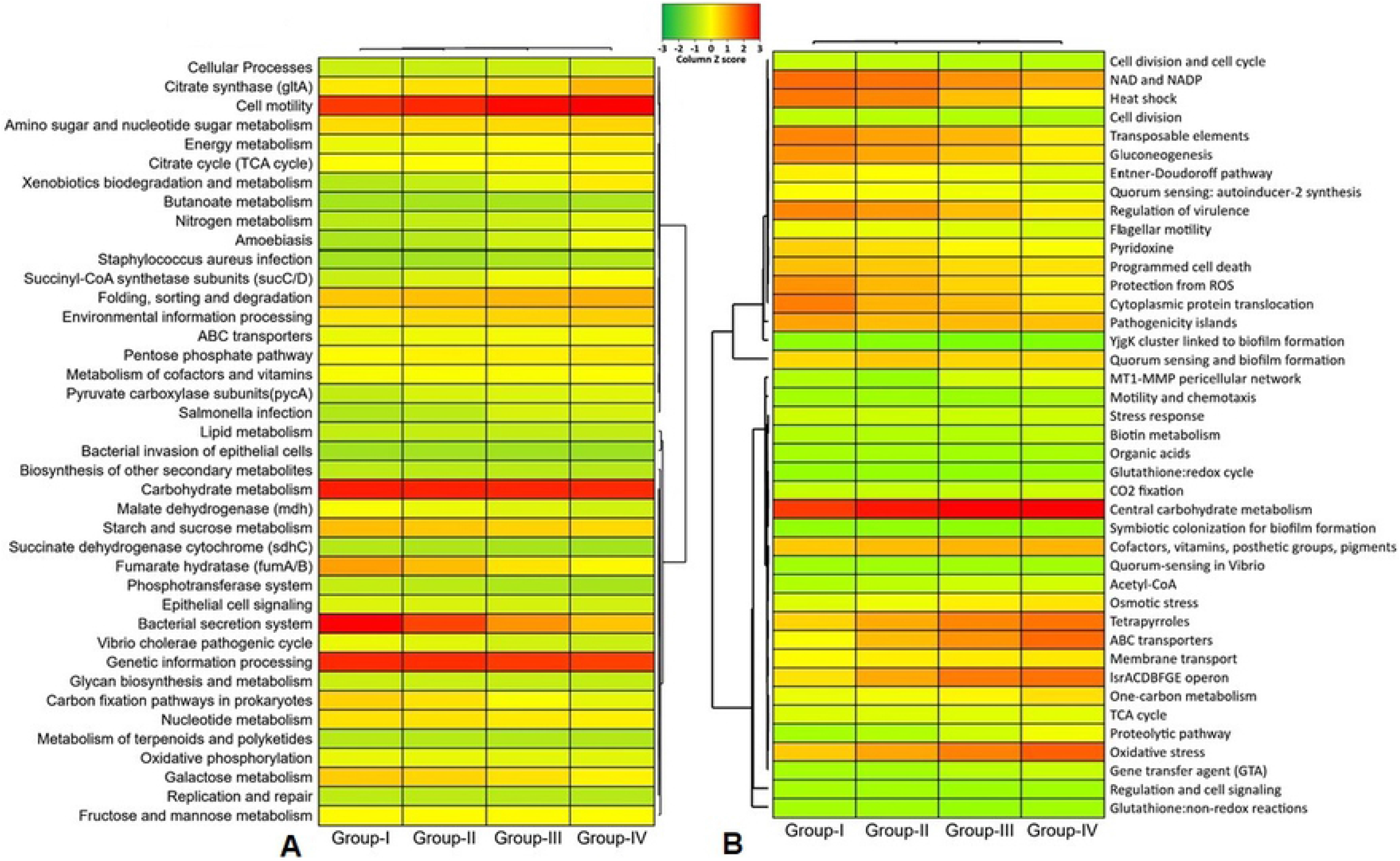
Functional genomic potentials of the anaerobic digestion associated microbial community through KEGG and SEED Pathways analysis. (A) Heatmap depicting the distribution of the 40 genes associated with the identified metabolic functional potentials detected by KEGG Pathways analysis within the four metagenomic groups of the AD microbiome. (B) Heatmap showing the distribution of the 41 functional gene composition and metabolic potential detected by SEED Pathways analysis within the four metagenomic groups of the AD microbiome. The color code indicates the presence and completeness of each gene, expressed as a value (Z score) between −2 (low abundance), and 2 (high abundance). The red color indicates the highest abundance whilst light green cells account for lower abundance of the respective genes in each metagenome.

We also found 41 statistically different (p = 0.033, Kruskal-Wallis test) SEED functions in the AD microbiomes. Overall, the top CH_4_ producing microbiomes (Group-III and Group-IV) had higher relative abundances of these SEED functions compared lower CH_4_ producing microbiomes (Group-I and Group-II), except for regulation of virulence (highest in Group-I microbes; 17.08%), gluconeogenesis (highest in Group-I microbes; 16.27%) and transposable elements (highest in Group-I microbes; 17.28%) (Fig 8B, Data S2). The Group-IV-microbiomes (highest CH_4_ producing) were enriched in genes coding for tetrapyrroles (17.42%), one carbon (10.29%) and biotin (4.55%) metabolism, oxidative (18.76%) and osmotic (9.94%) stress, proteolytic pathway (7.74%), MT1-MMP pericellular network (6.45%), acetyl-CoA production (5.33%) and motility and chemotaxis (3.13%) compared to the microbes of the other metagenomes. The Group-I microbiomes however had a higher abundance of SEED functions involved in protection from ROS (16.28%), heat shock (18.31%) and NAD and NADP (19.03%) (Fig 8B, Data S2).

## Discussion

This study is the first ever approach to reveal the dynamic shifts in microbiome composition and abundances in different levels of biogas production under the anaerobic digestion system using the state-of-the-art WMS technology along with analysis of the physicochemical parameters in Bangladesh. Anaerobic digestion of organic wastes is favored by the metabolic activities of different types of microorganisms including bacteria and archaea [32]. The physicochemical parameters assessment of the AD before and after digestion revealed that biogas production was in increasing trend up to Day 35 with periodic loading of slurry (Day 1, 2, 3, 8, 10, 14, 16, 24 and 35) at 1:1 ratio of raw cow dung and active sludge (Table S1). After 30 days of incubation, we found that cow wastes produced highest amount of biogas (on Day 35, CH_4_; 74.1%), and thereafter decrease gradually reaching 59.2% on Day 44 of processing (Fig 1). Biogas production chiefly depends on the content and chemical nature of biodegradable matter. The biochemical parameter of cow waste (slurry) reflects the presence of high content of readily biodegradable organic matter in the first phases (up to Day 35) of anaerobic digestion [32]. The CO_2_ concentration (%) found to be varied throughout the digestion process keeping an average value of 39.52%. Our analysis revealed that OC and TN content was higher at the time of loading of slurry in the AD compared to that of highest biogas production stage (at Day 35). The amount of carbon available of the substrate determines the maximum amount of CH_4_ and CO_2_ that can be formed by anaerobic digestion [7]. Conversely, C/N ratio remained lowest in this peak stage of CH_4_ production. Organic carbon is essential for bacterial growth, and determining of the C/N ratio is essential for optimal biogas production [45]. Moreover, the total content of phosphorus, Sulphur and heavy metals (chromium, lead and nickel) also remained lowest at this highest stage of biogas production (Day 35). Of note, the highest CH_4_ producing microbiomes were enriched with the higher relative abundance of genes coding for heavy metals (cobalt-zinc-cadmium, chromium compounds, arsenic, zinc, and cadmium) compared to the microbes of other three metagenomes. The lowest chromium, lead and nickel concentration during highest CH_4_ producing stage might be associated with their higher abundances of heavy metal resistance genes (Fig S6, Data S2), and small concentrations of these metals found in the process are essential for microbial maintenance [46,47]. Certain specific metals such as Sulphur, cobalt and nickel serve as cofactors in the enzymes involved in the formation of CH_4_ during anaerobic processing [48]. However, the minerals (e.g. zinc and copper) content of the AD did not vary throughout the digestion process revealing their important roles in various metabolic pathways of anaerobic digestion [46].

The within (alpha) and between (beta) sample diversity of the AD microbiomes showed that that microbial dysbiosis in the AD is closely linked to different levels of biogas production. Compared to loading phase of AD (Group-I), increased microbial diversity and species richness was observed in the later phases (Group-II, Group-III and Group-IV) of anaerobic digestion. Beta diversity also revealed a substantial microbial disparity in different levels of biogas production, and segregated the samples accordingly. Despite having higher taxonomic resolutions, the microbiomes of the AD remained inconsistent and fluctuates more in Group-II, Group-III and Group-IV than those of Group-I metagenome. The taxonomic annotations of the four groups of AD showed that they were a reservoir of bacteria, followed by archaea and viruses, which corroborated the findings of other studies [49,50]. Among the identified domains, bacteria dominated in abundance, comprising 81.80% of the total microbial populations, followed by archaea (15.43%), while viruses (2.77%) comprised the least abundant population. The observed high bacterial abundances suggest their crucial metabolic roles in biomass conversion and other reactions within the reactor systems [50]. The identified affiliates of archaea were mostly consumers of smaller substrates that were generated by the bacterial taxa. The archaeal species are able to use different methanogenic routes to convert the substrates into methane gas. Nevertheless, the main roles of the identified less abundant bacteriophages were unclear, though the strains could have been active in degrading other microbial cells in the AD systems [49,50].

In this study, *Firmicutes*, *Bacteroidetes*, *Proteobacteria*, *Actinobacteria*, *Spirochaetes* and *Fibrobacteres* were the most abundant bacterial phyla, and their relative abundance also varied according to the level of energy production in the corresponding sample groups. The observed community composition at phylum level, with dominance of *Firmicutes, Bacteroidetes* and *Proteobacteria*, is in line with previous findings for biogas reactors [47]. The first three phases (hydrolysis/cellulolysis, acidogenesis/fermentation and acetogenesis) of the anaerobic digestion are solely performed by fermentative Bacteria. In this study, the bacterial phyla *Firmicutes* (37.0% 42.0%) and *Bacteroidetes* (22.0% - 38.0%) appeared to dominate biogas communities in varying abundances depending on the apparent process conditions. The initial phase of the AD digestion process involves hydrolytic reactions that convert large macromolecules into smaller substrates [49]. In addition, the nature and composition of the substrates, availability of nutrients and ammonium/ammonia contents can affect both the composition and diversity of the methanogenic archaea [49]. Only certain methanogenic archaea are able to synthesize CH_4_ from the end products of bacterial fermentation. The performance and efficiency of these processes depend to a large extent on the presence of appropriate and adequate microorganisms along with the physicochemical conditions of the digester and quality of the substrates (organic materials) etc. The degree and rate of degradation (hydrolysis, fermentation, acetogenesis, and methanogenesis) and the biogas yield depend not only on the chemical and physical characteristics of the substrates, but also on the chosen process parameter such as temperature, humidity and retention time, that shape the composition of different microbial groups and communities that active in the process (Schnürer, 2016).

During the anaerobic process *Clostridiales* and *Bacteroidales* were identified as the top abundant order in all of the four metagenome groups. The syntrophic bacterial genera and strains of *Clostridiales order* are consumers of 30% of the generated electrons, which bypasses the rate limiting steps of the volatile fatty acids accumulation and contributes to AD stabilities [49,51]. In anaerobic environments, *Clostridiales* has been reported as the main cellulose degrader [52], and play important role in the hydrolysis step [47]. These findings are in line with many of the previous reports [49,53] who reported the potential roles of these bacterial taxa in the efficient and increased production of biogas under anaerobic condition [54]. In addition, bacteria belonging to the order *Bacteroidales* have been suggested to be involved in the degradation of lignocellulose materials, such as straw and hay, the chief component of cattle feed [54,55]. The identified affiliates of *Bacteroidales* belong to *Bacteroidetes*, and are majorly known to ferment carbohydrates and proteins, concomitantly releasing H_2_ [46]. Moreover, *Bacteroides*, the predominating genus in the AD coming from *Bacteroidales* order were observed to co-exist with methanogenic archaea possibly to increase energy extraction from indigestible plant materials [46], and we hypothesize that they are the key drivers of the observed *β*-diversities since the relative abundance of this genus gradually decreased with the digestion process.

The digestion process of the AD was carried out by the integrated cross-kingdom interactions since both bacteria and archaea were simultaneously detected in this WMS-based study corroborating with several earlier reports [25,26,28]. The strain-level taxonomic profiling revealed that methanogenic archaeal strains were more prevalent than bacterial strains. The archaeal domain of the AD microbiomes was composed of different strains of methanogenic, hydrogenotrophic and thermophilic genera of *Methanoculleus*, *Methanosarcina*, *Methanothrix*, *Methanobacterium* and *Methanobrevibacter* genera. The current findings are corroborated with many of the earlier studies who reported that these genera to be predominantly abundant in the AD of manures [32], and associated with biogas production under anaerobic conditions [56]. These methanogenic genera might reside in the microenvironments appropriate for anaerobic metabolism [28] and their presence has been reported in microbial communities producing biogas [57]. Members of *Methanoculleus* are hydrogenotrophic methanogens [58], while *Methanosarcina* species or strains are mostly acetoclasic but also able to use H_2_ [12,59]. In addition, *Methanosarcina* spp. has been reported to have higher growth rates and tolerance to pH changes and could potentially lead to stable methanogenesis in the AD [8,60]. *Methanobrevibacter* was predominant in initial phase of digestion (Day 2 and 15) in the bioreactor, and are known to be hydrogenotrophic, by using CO_2_ and H_2_ as substrates to generate biomethane [12,57]. These archaeal genera are suggested to play vital role in hydrogenotrophic methanogenesis, and maintaining methanogenic community diversity [24,27].

Indicator species (IS) and network analyses were used to identify at higher taxonomic resolution of the individual bacterial and/or archaeal species with enrichment or depletion patterns in different phases of anaerobic digestion. These analyses revealed the association of different methanogenic species with increased level of CH_4_ production [61], and all were observed to be more abundant in highest CH_4_ producing metagenome (Group-IV). However, common indicator bacteria *E. coli*, *Salmonella* and *Staphylococcus* species were not found in indicator species analysis, and these findings are supported by several previous reports of the absence of common indicator bacteria after 30 days digestion in the experimental AD at different temperatures (25 °C – 45 °C) [62]. Spearmen correlation analysis showed negative correlation between Group-II and Group-IV microbiomes. These findings are in line with the metabolic functional potentials of the AD microbiomes since fumarate hydratase (*fumA*/B), malate dehydrogenase (*mdh*) and bacterial secretion system associated genes were predominantly overexpressed in Group-I and Group-III microbiomes, which gradually decreased with advance of digestion process, and remained more than two-fold lower expressed in highest CH_4_ production stage (Group-IV) (Fig 8A, Data S2). The microbial communities present in the early phage of digestion process (Group-I) increased both composition and abundances in the second phase of digestion (Group-II metagenome) in ambient growth conditions of the AD. With the advance of digestion time, the increasing efficiency of anaerobic digestion creates a favorable environment for the methanogens (Figs 4 and 6), and thus found in higher composition and abundances in Group-III and Group-IV metagenomes [24,27]. In addition, the Group-IV-microbiomes showed higher genomic functional activities related to tetrapyrroles, one carbon and biotin metabolism, oxidative and osmotic stress, proteolytic pathways, MT1-MMP pericellular network, acetyl-CoA production, and motility and chemotaxis compared to the microbes of the other groups. Though, these findings also support the taxonomic dysbiosis of microbiomes in Group-IV metagenome, however further comprehensive study is needed to elucidate the modulation of microbiome shifts, their functional potentials and genomic expression using a larger dataset.

## Conclusions

The level of biogas production increased gradually up to Day 35 (highest CH_4_ concentration), and declined thereafter under controlled environment of the AD. With the increase of CH_4_ production, the amount of non-metallic elements (phosphorus and Sulphur) and heavy metals (chromium, lead and nickel) decreases at Day 35 of the digestion process. The pair-wise comparison of the within (alpha) and between (beta) sample diversities revealed that the microbiomes of the Group-IV significantly differed with those of other three groups. The present study revealed an imbalance distribution of bacterial phyla across four groups keeping comparatively higher relative abundances and compositions of the methanogenic microbiomes during highest CH_4_ producing stage (Group-IV). The indicator species analysis revealed that Group-III and Group-IV were highly specific for energy production. The correlation network analysis of the indicator species showed that different strains of *Euryarcheota* and *Firmicutes* phyla were negatively correlated but associated with highest level of energy production.

## Acknowledgements

This project is financed by the Surge Engineering (www.surgeengineering.com) Bangladesh.

## Supplementary Information

Supplementary information supporting the results of the study are available in this article as Data S1 and S2, Figs. S1-S6, and Table S1.

## Author Contributions

MNH performed bioinformatics analysis, visualized figures, interpreted results and drafted the original manuscript. MSR carryout field experiment, curated the data and performed bioinformatics analysis. JAP carryout field experiment, and physicochemical analysis. MRI and MAS edited the drafted manuscript. ND, MAH and MS conceived the study and critically reviewed the drafted manuscript.

## Competing Interests

The authors declare no competing interests.

## Data availability

The sequence data reported in this article have also been deposited in the National Center for Biotechnology Information (NCBI) under BioProject accession number PRJNA668799.

## Ethics Statement

Not applicable

## Notes

### Competing Interest Statement

The authors have declared no competing interest.

## References

1. Khan EU, Mainali B, Martin A, Silveira S (2014) Techno-economic analysis of small scale biogas based polygeneration systems: Bangladesh case study. Sustainable Energy Technologies and Assessments 7: 68–78.

2. Khan EU, Martin AR (2016) Review of biogas digester technology in rural Bangladesh. Renewable and Sustainable Energy Reviews 62: 247–259.

3. Weiland P (2010) Biogas production: current state and perspectives. Applied microbiology and biotechnology 85: 849–860.

4. Wirth R, Kovács E, Maróti G, Bagi Z, Rákhely G, et al. (2012) Characterization of a biogas-producing microbial community by short-read next generation DNA sequencing. Biotechnology for biofuels 5: 1–16.

5. Das A, Sahoo S, Rana S (2018) Sustainable conservation of kitchen wastes into fuels and organic fertilizer. International Journal of Engineering Sciences and Research Technology 7: 503–510.

6. Agostini A, Battini F, Padella M, Giuntoli J, Baxter D, et al. (2016) Economics of GHG emissions mitigation via biogas production from Sorghum, maize and dairy farm manure digestion in the Po valley. Biomass and Bioenergy 89: 58–66.

7. Mulka R, Szulczewski W, Szlachta J, Prask H (2016) The influence of carbon content in the mixture of substrates on methane production. Clean Technologies and Environmental Policy 18: 807–815.

8. Tian Z, Cabrol L, Ruiz-Filippi G, Pullammanappallil P (2014) Microbial ecology in anaerobic digestion at agitated and non-agitated conditions. PLOS one 9: e109769.

9. Campanaro S, Treu L, Rodriguez-R LM, Kovalovszki A, Ziels RM, et al. (2020) New insights from the biogas microbiome by comprehensive genome-resolved metagenomics of nearly 1600 species originating from multiple anaerobic digesters. Biotechnology for biofuels 13: 1–18.

10. Valentinuzzi F, Cavani L, Porfido C, Terzano R, Pii Y, et al. (2020) The fertilising potential of manure-based biogas fermentation residues: Pelleted vs. liquid digestate. Heliyon 6: e03325.

11. Fernandez-Gonzalez N, Braz G, Regueiro L, Lema J, Carballa M (2020) Microbial invasions in sludge anaerobic digesters. Applied Microbiology and Biotechnology: 1–13.

12. Zhu X, Campanaro S, Treu L, Seshadri R, Ivanova N, et al. (2020) Metabolic dependencies govern microbial syntrophies during methanogenesis in an anaerobic digestion ecosystem. Microbiome 8: 1–14.

13. Theuerl S, Klang J, Hülsemann B, Mächtig T, Hassa J (2020) Microbiome Diversity and Community-Level Change Points within Manure-based small Biogas Plants. Microorganisms 8: 1169.

14. Ziels RM, Svensson BH, Sundberg C, Larsson M, Karlsson A, et al. (2018) Microbial rRNA gene expression and co-occurrence profiles associate with biokinetics and elemental composition in full-scale anaerobic digesters. Microbial biotechnology 11: 694–709.

15. Mukti RF, Sinthee SS (2019) Metagenomic Approach: Transforming In-Silico Research for Improved Biogas Production. International Journal of Applied Sciences and Biotechnology 7: 6–11.

16. Kemausuor F, Adaramola MS, Morken J (2018) A review of commercial biogas systems and lessons for Africa. Energies 11: 2984.

17. Bremges A, Maus I, Belmann P, Eikmeyer F, Winkler A, et al. (2015) Deeply sequenced metagenome and metatranscriptome of a biogas-producing microbial community from an agricultural production-scale biogas plant. Gigascience 4: s13742-13015-10073-13746.

18. Ziganshin AM, Ziganshina EE, Kleinsteuber S, Nikolausz M (2016) Comparative analysis of methanogenic communities in different laboratory-scale anaerobic digesters. Archaea 2016.

19. Pham JV, Yilma MA, Feliz A, Majid MT, Maffetone N, et al. (2019) A review of the microbial production of bioactive natural products and biologics. Frontiers in microbiology 10: 1404.

20. Sundberg C, Al-Soud WA, Larsson M, Alm E, Yekta SS, et al. (2013) 454 pyrosequencing analyses of bacterial and archaeal richness in 21 full-scale biogas digesters. FEMS microbiology ecology 85: 612–626.

21. Hahnke S, Langer T, Klocke M (2018) Proteiniborus indolifex sp. nov., isolated from a thermophilic industrial-scale biogas plant. International journal of systematic and evolutionary microbiology 68: 824–828.

22. Hoque M, Das Z, Rahman A, Haider M, Islam M (2018) Molecular characterization of Staphylococcus aureus strains in bovine mastitis milk in Bangladesh. International journal of veterinary science and medicine 6: 53–60.

23. Saha O, Hoque MN, Islam OK, Rahaman M, Sultana M, et al. (2020) Multidrug-resistant avian pathogenic Escherichia coli strains and association of their virulence genes in Bangladesh. Microorganisms 8: 1135.

24. Zhang Q, Wang M, Ma X, Gao Q, Wang T, et al. (2019) High variations of methanogenic microorganisms drive full-scale anaerobic digestion process. Environment international 126: 543–551.

25. Hoque MN, Istiaq A, Clement RA, Gibson KM, Saha O, et al. (2020) Insights into the resistome of bovine clinical mastitis microbiome, a key factor in disease complication. Frontiers in Microbiology 11: 860.

26. Hoque MN, Istiaq A, Clement RA, Sultana M, Crandall KA, et al. (2019) Metagenomic deep sequencing reveals association of microbiome signature with functional biases in bovine mastitis. Scientific reports 9: 1–14.

27. De Vrieze J, Christiaens ME, Walraedt D, Devooght A, Ijaz UZ, et al. (2017) Microbial community redundancy in anaerobic digestion drives process recovery after salinity exposure. Water research 111: 109–117.

28. Hoque MN, Istiaq A, Rahman MS, Islam MR, Anwar A, et al. (2020) Microbiome dynamics and genomic determinants of bovine mastitis. Genomics 112: 5188–5203.

29. Sparks DL, Page A, Helmke P, Loeppert RH (2020) Methods of soil analysis, part 3: Chemical methods: John Wiley & Sons.

30. Afilal M, Elasri O, Merzak Z (2014) Caractérisations des déchets organiques et évaluation du potentiel Biogaz (Organic waste characterization and evaluation of its potential biogas). J Mater Environ Sci 5: 2014.

31. Banerjee P, Prasad B (2020) Determination of concentration of total sodium and potassium in surface and ground water using a flame photometer. Applied Water Science 10: 1–7.

32. El Asri O, Afilal ME, Laiche H, Elfarh A (2020) Evaluation of physicochemical, microbiological, and energetic characteristics of four agricultural wastes for use in the production of green energy in Moroccan farms. Chemical and Biological Technologies in Agriculture 7: 1–11.

33. Hong C, Manimaran S, Shen Y, Perez-Rogers JF, Byrd AL, et al. (2014) PathoScope 2.0: a complete computational framework for strain identification in environmental or clinical sequencing samples. Microbiome 2: 1–15.

34. Glass EM, Wilkening J, Wilke A, Antonopoulos D, Meyer F (2010) Using the metagenomics RAST server (MG-RAST) for analyzing shotgun metagenomes. Cold Spring Harbor Protocols 2010: pdb.prot5368.

35. Wood DE, Lu J, Langmead B (2019) Improved metagenomic analysis with Kraken 2. Genome biology 20: 257.

36. Li H (2018) Minimap2: pairwise alignment for nucleotide sequences. Bioinformatics 34: 3094–3100.

37. Li H, Durbin R (2010) Fast and accurate long-read alignment with Burrows–Wheeler transform. Bioinformatics 26: 589–595.

38. Li H, Handsaker B, Wysoker A, Fennell T, Ruan J, et al. (2009) The sequence alignment/map format and SAMtools. Bioinformatics 25: 2078–2079.

39. Koh H (2018) An adaptive microbiome α-diversity-based association analysis method. Scientific reports 8: 1–12.

40. Beck J, Holloway JD, Schwanghart W (2013) Undersampling and the measurement of beta diversity. Methods in Ecology and Evolution 4: 370–382.

41. McMurdie PJ, Holmes S (2013) phyloseq: an R package for reproducible interactive analysis and graphics of microbiome census data. PloS one 8: e61217.

42. De Cáceres M, Legendre P, Wiser SK, Brotons L (2012) Using species combinations in indicator value analyses. Methods in Ecology and Evolution 3: 973–982.

43. Doster E, Lakin SM, Dean CJ, Wolfe C, Young JG, et al. (2020) MEGARes 2.0: a database for classification of antimicrobial drug, biocide and metal resistance determinants in metagenomic sequence data. Nucleic acids research 48: D561–D569.

44. Kanehisa M, Sato Y, Furumichi M, Morishima K, Tanabe M (2019) New approach for understanding genome variations in KEGG. Nucleic acids research 47: D590–D595.

45. Tanimu MI, Ghazi TIM, Harun MR, Idris A (2015) Effects of feedstock carbon to nitrogen ratio and organic loading on foaming potential in mesophilic food waste anaerobic digestion. Applied microbiology and biotechnology 99: 4509–4520.

46. Agustini CB, da Costa M, Gutterres M (2020) Biogas from Tannery Solid Waste Anaerobic Digestion Is Driven by the Association of the Bacterial Order Bacteroidales and Archaeal Family Methanosaetaceae. Applied biochemistry and biotechnology: 1–12.

47. Liu T, Sun L, Müller B, Schnürer A (2017) Importance of inoculum source and initial community structure for biogas production from agricultural substrates. Bioresource technology 245: 768–777.

48. Zandvoort MH, van Hullebusch ED, Gieteling J, Lens PN (2006) Granular sludge in full-scale anaerobic bioreactors: trace element content and deficiencies. Enzyme and microbial technology 39: 337–346.

49. Muturi SM, Muthui LW, Njogu PM, Onguso JMa, Wachira FN, et al. (2021) Metagenomics survey unravels diversity of biogas microbiomes with potential to enhance productivity in Kenya. Plos one 16: e0244755.

50. Stolze Y, Zakrzewski M, Maus I, Eikmeyer F, Jaenicke S, et al. (2015) Comparative metagenomics of biogas-producing microbial communities from production-scale biogas plants operating under wet or dry fermentation conditions. Biotechnology for biofuels 8: 1–18.

51. Schmidt A, Müller N, Schink B, Schleheck D (2013) A proteomic view at the biochemistry of syntrophic butyrate oxidation in Syntrophomonas wolfei. PloS one 8: e56905.

52. Lynd LR, Weimer PJ, Van Zyl WH, Pretorius IS (2002) Microbial cellulose utilization: fundamentals and biotechnology. Microbiology and molecular biology reviews 66: 506–577.

53. Luo G, Fotidis IA, Angelidaki I (2016) Comparative analysis of taxonomic, functional, and metabolic patterns of microbiomes from 14 full-scale biogas reactors by metagenomic sequencing and radioisotopic analysis. Biotechnology for biofuels 9: 1–12.

54. Sun L, Pope PB, Eijsink VG, Schnürer A (2015) Characterization of microbial community structure during continuous anaerobic digestion of straw and cow manure. Microbial biotechnology 8: 815–827.

55. Hanreich A, Schimpf U, Zakrzewski M, Schlüter A, Benndorf D, et al. (2013) Metagenome and metaproteome analyses of microbial communities in mesophilic biogas-producing anaerobic batch fermentations indicate concerted plant carbohydrate degradation. Systematic and applied microbiology 36: 330–338.

56. Kouzuma A, Tsutsumi M, Ishii Si, Ueno Y, Abe T, et al. (2017) Non-autotrophic methanogens dominate in anaerobic digesters. Scientific reports 7: 1–13.

57. Poehlein A, Schneider D, Soh M, Daniel R, Seedorf H (2018) Comparative genomic analysis of members of the genera Methanosphaera and Methanobrevibacter reveals distinct clades with specific potential metabolic functions. Archaea 2018.

58. Shcherbakova V, Rivkina E, Pecheritsyna S, Laurinavichius K, Suzina N, et al. (2011) Methanobacterium arcticum sp. nov., a methanogenic archaeon from Holocene Arctic permafrost. International journal of systematic and evolutionary microbiology 61: 144–147.

59. Kor-Bicakci G, Ubay-Cokgor E, Eskicioglu C (2020) Comparative analysis of bacterial and archaeal community structure in microwave pretreated thermophilic and mesophilic anaerobic digesters utilizing mixed sludge under organic overloading. Water 12: 887.

60. Cho S-K, Im W-T, Kim D-H, Kim M-H, Shin H-S, et al. (2013) Dry anaerobic digestion of food waste under mesophilic conditions: performance and methanogenic community analysis. Bioresource Technology 131: 210–217.

61. Deng S, Wipf HM-L, Pierroz G, Raab TK, Khanna R, et al. (2019) A plant growth-promoting microbial soil amendment dynamically alters the strawberry root bacterial microbiome. Scientific reports 9: 1–15.

62. Dinova N, Belouhova M, Schneider I, Rangelov J, Topalova Y (2018) Control of biogas production process by enzymatic and fluorescent image analysis. Biotechnology & Biotechnological Equipment 32: 366–375.

